# Passive symplastic phloem loading in the duckweed *Spirodela polyrhiza*

**DOI:** 10.1101/2025.01.27.635011

**Authors:** Jiazhou Li, Johannes Liesche

## Abstract

The lemnoidae, commonly called duckweeds, are a group of small, rapidly growing aquatic plants that play an important role in pond ecosystems and are used in biotechnological and remediation applications. While small, duckweeds feature phloem tissue in fronds and roots. To gain insight on duckweed phloem function, we investigated how sugar is loaded into the phloem sieve elements in the giant duckweed *Spirodela polyrhiza*. Genomes of *S. polyrhiza* and three other duckweeds do not feature genes for the sucrose transporters typically associated with active apoplastic phloem loading. Neither did a sucrose transporter inhibitor affect sucrose concentration in phloem exudate. Active symplastic phloem loading was excluded based on the conventional plasmodesmata configuration and absence of oligosaccharides in *S. polyrhiza* phloem. Instead, uniform plasmodesmata density along the phloem loading pathway indicated a passive symplastic phloem loading type. When plasmodesmata permeability was artificially reduced by hormone treatment, the sucrose concentration in the phloem exudate was reduced, highlighting the potential role of plasmodesmata regulation in setting carbon export rates in species with passive phloem loading. Our results identify *S. polyrhiza* as the first monocot species with passive phloem loading. Moreover, they indicate opportunities for optimization of duckweed growth.

## Introduction

Vascular plants are generally assumed to depend on sugar transport in the phloem for an efficient distribution of carbon and energy between organs. However, in how far this applies to the small aquatic plants of the lemnoidae (duckweeds) has not been investigated. The role of xylem transport differs strongly from other plants, as it is not required for nutrient transport from roots to fronds (Ware et al. 2023). Indeed, the minimal impact that root removal had on the frond area growth rate of many duckweed species (Ware et al. 2023), suggests that phloem function might be different, too. However, duckweeds do have phloem tissues with similar structure to their monocot relatives on land (Melaragno and Walsh 1976; Kim 2007). Besides supporting root development and growth, phloem transport could facilitate carbon storage in the root, or sustain sugar secretion from the roots for shaping the rhizosphere microbiome (Acosta et al. 2020; Loo et al. 2024).

A key step of phloem transport is the loading of sugars produced by photosynthesis in source tissues into the phloem sieve elements. Plants employ three primary strategies for phloem loading: active apoplastic, active symplastic, and passive symplastic (Liesche and Patrick 2017). Active apoplastic loading involves the energy-dependent transport of sucrose into the phloem companion cell-sieve element complex by membrane transporters. All monocots analyzed so far, which are all grass species, are active apoplastic loaders (Zhang and Turgeon 2018). Active symplastic loading, also called polymer trapping, is an energy-dependent process, too. Sucrose is converted into oligosaccharides within companion cells, which are then transported in the sieve elements (Rennie & Turgeon, 2009). In contrast, passive symplastic loading relies on the concentration gradient to drive sucrose diffusion from mesophyll cells to the companion cell-sieve element- complex through plasmodesmata (Liesche 2017).

Investigating the phloem loading strategy of duckweeds could yield important insights into the carbon-use-efficiency and adaptability of these small aquatic plants. Duckweeds are used in approaches for wastewater treatment, biofuel production, and as a model organism for understanding plant biology and ecology in aquatic environments (Cui and Cheng 2014; Chen et al., 2020; Zhang et al., 2021). Our study focuses on *Spirodela polyrhiza*, the ‘greater duckweed’. This species is known for its larger size compared to other duckweed species, with a frond diameter of 0.5 cm to 1 cm and multiple roots of about 2 cm to 4 cm length.

## Materials and Methods

### Plant material and growth conditions

*Spirodela polyrhiza* was cultivated for 2–3 weeks in 150 mL of sterile half-strength Shenk-Hildebrandt (1/2 SH) medium supplemented with 1% (w/v) sucrose, adjusted to pH 5.6, as described by Schenk and Hildebrandt (1972). The culture was maintained in a light incubator (HYG-1050D-4, HuaYi Instrument) under a light intensity of 90 μmol photons m^−2^ s^−1^. A 16-hour light/8-hour dark photoperiod was used, with day temperatures set at 22°C and night temperatures at 20°C.

Hormone treatment of *S. polyrhiza* was carried out on growth medium containing 100 μM ABA (Solarbio Science & Technology, Beijing, China), 100 μM SA (Solarbio Science & Technology, Beijing, China), 10 μM GA (Solarbio Science & Technology, Beijing, China), and 10 μM NAA (Solarbio Science & Technology, Beijing, China), respectively. *S. polyrhiza* plants with similar size and morphology were transferred to the treatment media and cultivated for 12 hours under conditions described above.

### Phylogenetic analysis

The genomic data for *S. polyrhiza* were retrieved from Phytozome13 (https://phytozome-next.jgi.doe.gov/), including the genome file (Spolyrhiza_290_v1.fasta) and the gene annotation file (Spolyrhiza_290_v2.gene.gff3). Gene family identification and analysis were conducted using TB-tools (Chen et al., 2020). Protein sequences of the SUT and SWEET families from nine species, including *Arabidopsis thaliana*, *Solanum lycopersicum*, *Solanum tuberosum*, *Glycine max*, *Populus tremula x Populus alba*, *Zea mays*, *Hordeum vulgare*, *Triticum aestivum*, *Lemna minor 7210*, *Lemna japonica 9421*, *Wolffia australiana 8730*, and O*ryza sativa*, were retrieved from the NCBI GeneBank and JGI database (see Table S1 and Table S2). All sequences were converted into FASTA format and compiled into a single file.

Sequence alignment and phylogenetic tree construction were conducted as described by Chen et al. (2020), using the nearest neighbor clustering method implemented in MEGA7 software (Kumar et al., 2016).

### Yeast complementation assays

Suitable restriction enzyme sites were chosen to insert the coding sequences of *SpSUT2*, *SpSWEET1*, *SpSWEET11*, and *AtSUC2* into the vector pDR196. Details of the restriction enzymes and primers used for amplification are provided in Supplemental Table S3 and Table S4. Yeast was cultured in YPDA medium, and the genetic transformation process was carried out according to Liu et al. (2022). Yeast drop test assays involving the mutant strains *EBY.VW4000* or *SUSY7/ura3* were conducted following the method described by Wang et al. (2020) on SD/ura medium with 3-5 days between inoculation and analysis. The yeast strain *EBY.VW4000* was employed to assess the glucose transport activity of the duckweed transporter proteins, while the *SUSY7/ura3* mutant strain was used to evaluate their ability to facilitate sucrose transport.

### *In situ* hybridization

Tissue samples were fixed in 4% paraformaldehyde in a neutral buffer (PBS, pH 7) to preserve structural integrity and nucleic acids (Tautz & Pfeifle, 1989). After fixation, tissues were cryo-sectioned with 10 µm section thickness and mounted onto poly-L-lysine-coated slides (Murray & Barnes, 2012). Sections were pretreated to enhance probe access, including permeabilization and blocking of nonspecific binding (Liu et al., 2014). RNA probes (Table S5), labeled with digoxigenin (DIG), were synthesized based on target sequences. For hybridization, sections were incubated with a prehybridization solution (Boster Biological Technology, WuHan, China) to block nonspecific binding (Müller et al., 2010). DIG-labeled probes were applied and incubated at the optimal temperature of 55 °C for hybridization (Frohlich & Majewski, 2016). Post-hybridization, sections were washed stringently to remove unbound probes (Tautz & Pfeifle, 1989). Hybridized probes were detected using anti-DIG antibodies conjugated to alkaline phosphatase, with signal visualization using a colorimetric system (Pannoramic 250Flash, 3DHISTECH, Hungary) (Müller et al., 2010). Slides were mounted and analyzed under a microscope (DM2000 LED, Leicia).

### Sugar concentration measurements

Sugars from duckweed fronds were extracted using ethanol. The fronds were harvested, weighed, and placed into a 2 mL centrifuge tube containing small steel beads. To maintain sample freshness, the fronds were immediately frozen in liquid nitrogen. They were then ground into a fine powder using a high-throughput tissue grinder. Afterwards, 1 mL of 80% ethanol was added to the centrifuge tube, which was incubated in a water bath at 80°C for 30 minutes to facilitate thorough sugar extraction. The mixture was centrifuged at 12,000 × g for 20 minutes, and the supernatant was collected. The extraction was repeated to ensure complete extraction, and all supernatants were combined. The supernatant was concentrated using a vacuum concentrator at 60°C to remove residual ethanol. The remaining solid material was suspended in 2 mL of ddH_2_O. To purify the sample, the solution was filtered through a 0.22 µm water-phase micropore filter and rapidly frozen in liquid nitrogen. The final samples were stored at -80°C for subsequent ion chromatography analysis.

Phloem exudate was collected using the EDTA-facilitated exudation method (Xu et al., 2018). Freshly excised fronds were immersed in 10 mM EDTA solution (pH 7.5) to induce exudation, and the sap was collected and stored at −20°C. Sugar concentrations were measured following the method described by Xu et al. (2018) using an ion chromatograph (ICS-5000+, Thermo Fisher, CA, USA) equipped with an integrated pulsed amperometric detector. A CarboPac PA1 chromatographic column (4 × 250 mm i.d., 10 µm) served as the separation column, while 2 M NaOH was utilized as the mobile phase at a flow rate of 1 mL/min. Soluble sugar concentrations were quantified by normalizing peak areas, with data analysis performed using Chromeleon 7 software (Thermo Fisher, CA, USA). Note that the EDTA-facilitated exudation method is not suitable for absolute quantification of phloem sap sugars (Xu et al. 2019), meaning that values cannot be directly compared to whole frond sugar measurements.

The effect of different concentrations of pCMBS was evaluated by preparing 1 mM and 0.5 mM solutions in sterile water. Prior to the application of pCMBS a 0.1% (w/v) Tween-20 solution was sprayed as a pretreatment to enhance the absorption of pCMBS by plant leaves. Treatments were applied at the transition from day to night, with both concentrations of pCMBS sprayed onto duckweed fronds, and ddH_2_O used as the control.

### Transmission electron microscopy

Squares of approximately 5 mm width were excised from mature fronds and immediately fixed in 2.5% glutaraldehyde in 0.1 M PBS (pH 7.2) for a minimum of 12 hours at 4°C. The samples were then rinsed four times in fresh 0.1 M PBS (pH 7.2) for 15 minutes each, followed by incubation in 1% osmium tetroxide at 4°C for 4 hours to enhance contrast. Subsequently, the samples were washed in phosphate buffer and dehydrated through a graded ethanol and acetone series. The tissue was embedded in LR White resin and sectioned to 70 nm thickness using an ultramicrotome (EM UC7, Leica Microsystems). Sections were stained sequentially with 2% uranyl acetate for 20 minutes, followed by lead citrate for 15 minutes to enhance electron contrast. Finally, the prepared samples were observed and imaged using a transmission electron microscope (HT7800, HITACHI, Tokyo, Japan). PD density was determined from electron micrographs as described by Liesche et al. (2019), meaning that PDs were counted at the different interfaces per minor vein and counts multiplied to indicate numbers along 1 µm of minor vein.

### Measurement of plasmodesmata-mediated cell wall permeability

*S. polyrhiza* fronds were used in Fluorescence Recovery After Photobleaching (FRAP) experiments. The cytosolic tracer carboxyfluorescein diacetate (cFDA, ThermoFisher, Waltham, MA, USA) was prepared at a concentration of 50 µM in PBS buffer (pH 7.2). Following hormone treatment, whole plants were incubated in the cFDA solution for 20 minutes. After rinsing with PBS buffer, the leaves were placed on a glass slide and observed using a confocal microscope (Leica SP8, Leica Microsystems). A 20X objective lens was used, with fluorescence excitation at 488 nm, and emission detection at 494 to 524 nm. Once an appropriate region was identified, the system was switched to FRAP protocol. For bleaching, the 488 nm laser intensity to 100%, instead of 15% for imaging. An appropriate bleaching region was selected, and the bleaching time course was set as follows: pre-bleach for 2 frames, bleaching for 10 frames, post-bleach for 30 frames. Upon completion of the experiment, the average intensity values across all frames in the bleached regions were calculated. Non-linear regression analysis of the post-bleach data was performed using GraphPad Prism 8 (GraphPad Software, San Diego, CA, USA) to determine the half-time of recovery.

### Aniline Blue Staining

To observe the callose in the epidermal cells of *S. polyrhiza* leaves, we adapted the method of Zavaliev and Epel (2015) with some modifications. The experiment began by collecting intact *S. polyrhiza* fronds, which were then immersed in 96% ethanol for decolorization. After decolorization, the leaves were washed with distilled water to remove residual ethanol and rehydrated in distilled water containing 0.01%(w/v) Tween-20 for 1 hour. After rehydration, approximately 5 mm × 2 mm strips were cut from each leaf. The samples were then placed in a staining solution containing 0.1% (w/v) aniline blue and 0.01 M K_3_PO_4_ (pH 10.5) and subjected to vacuum treatment in a centrifuge concentrator (Concentrator Plus, Eppendorf, Germany) for 10 minutes to ensure that the staining solution entered the cells. Following staining, the samples were shaken at 40 rpm at room temperature for 1 hour to ensure even staining. The stained samples were then observed using a Leica SP8 confocal laser scanning microscope (Leica Microsystems, Germany), with 405 nm excitation and 475–525 nm emission detection settings. The pinhole was set to 2 Airy Units, and a 40x objective lens was used. After image acquisition, a circular region of interest with a diameter of 1.5 µm was drawn around each aniline blue spot using ImageJ (Schindelin et al., 2012), and the mean intensity was recorded.

### Transcriptomics

Total RNA was extracted using the TRIzol reagent (Invitrogen, CA, USA) according to the manufacturer’s protocol. RNA purity and quantification were evaluated using the NanoDrop 2000 spectrophotometer (Thermo Scientific, CA, USA). RNA integrity was assessed using the Agilent 2100 Bioanalyzer (Agilent Technologies, Santa Clara, CA, USA). The libraries were constructed using Universal V6 RNA-seq Library Prep Kit (Vazyme Biotech, NanJing, China) according to the manufacturer’s instructions. The libraries were sequenced on an llumina Novaseq 6000 platform (llumina, CA, USA) and 150 bp paired-end reads were generated. This analysis involved parametric transcriptome sequencing of 15 samples, yielding a total of 101.37 Gb of clean data. The effective data volume per sample ranged from 5.54 to 7.04 Gb, with Q30 base quality scores between 91.5% and 92.34%, and an average GC content of 55.88%. Reads were aligned to the reference genome, resulting in alignment rates ranging from 89.53% to 90.59%. The transcriptome sequencing was conducted by OE Biotech Co., Ltd. (Shanghai, China). The RNA-seq data of this study is available at the NCBI Sequence Read Achieve (http://www.ncbi.nlm.nih.gov/geo) with the accession number [will be added].

### Statistics

Fitting of the recovery curves in FRAP experiments to evaluate PD permeability was performed by nonlinear fitting in GraphPad software (GraphPad Software, San Diego, CA, USA). To evaluate the significance of differences between samples, pairwise Student’s *t*-tests were conducted. A level of significance of *P* < 0.05 was assumed. Sample numbers are provided in the figure legends.

## Results

### Duckweeds lack type IIB sucrose transporters

Our analysis of the SUT gene family in duckweeds, represented by *Spirodela polyrhiza*, *Lemna minor* 7210, *Lemna japonica* 9421, and *Wolffia australiana* 8730, revealed significant differences compared to other monocot species (Fig. 1). Duckweeds lacked Type IIB sucrose transporters (SUTs), which mediate active phloem loading in other monocots such as rice (*Oryza sativa*), maize (*Zea mays*), and barley (*Hordeum vulgare*) (Reinders et al. 2012; Slewinski et al., 2009). As expected, duckweed genomes did not feature Type I SUTs associated with active phloem loading in dicots (Fig. 1A, Reinders et al. 2012). Like all embryophytes, duckweeds possess Type IIA and Type IV SUTs, typically associated with sucrose transport in sink tissues and intracellular sucrose storage, respectively (Fig. 1A; Braun et al., 2014; Weise et al., 2000).

**Fig. 1.**
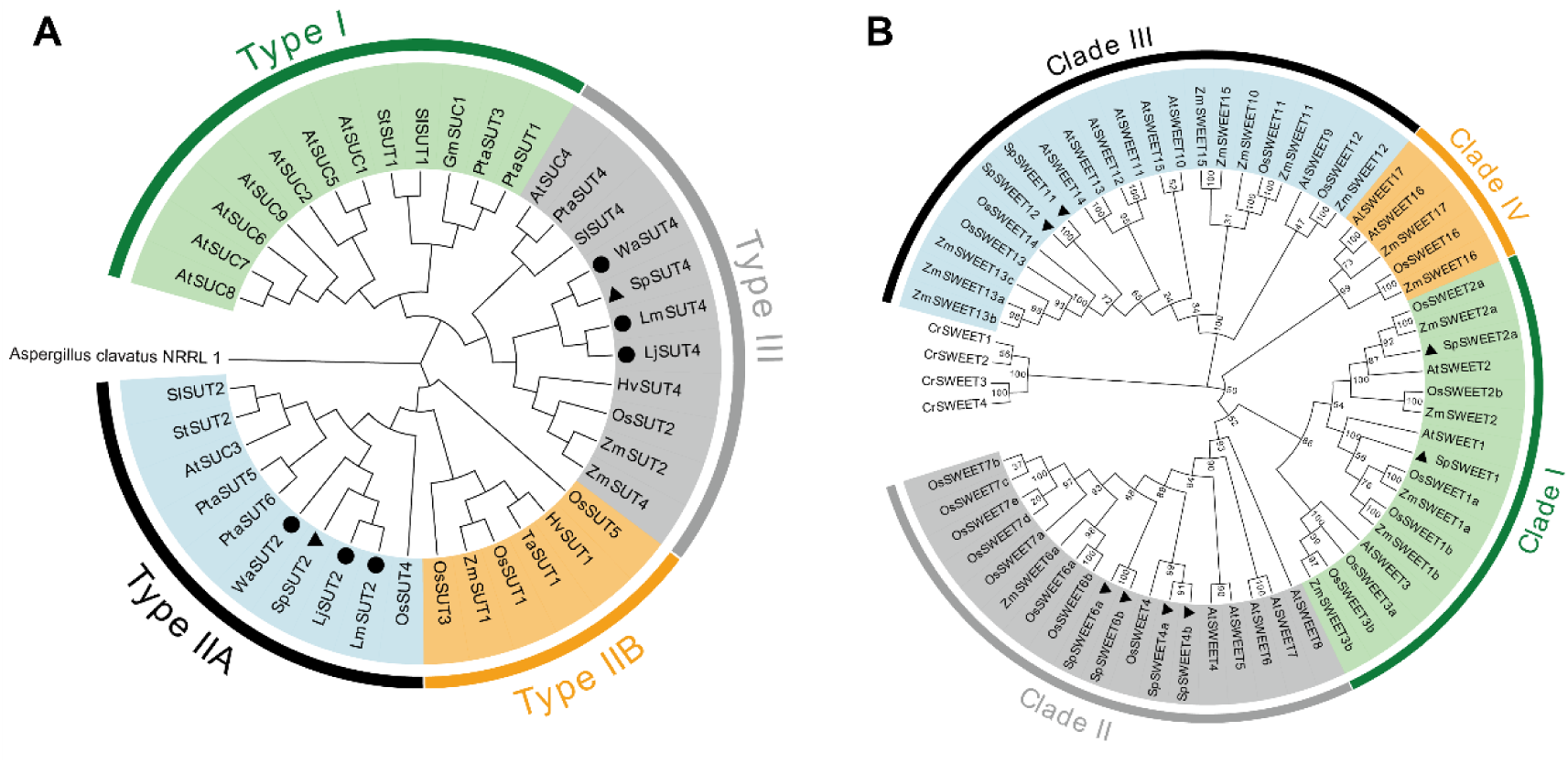
Phylogenetic analysis of *SUT* and *SWEET* gene families. Phylogenetic trees based on SUT (A) and SWEET (B) protein sequences. *Spirodela polyrhiza transporters* are marked by black triangles while the SUTs of other duckweeds are marked by filled circles. Species abbreviations: At – *Arabidopsis thaliana*, Cr –*Chlamydomonas reinhardtii*, Gm – *Glycine max (soybean)*, Hv – *Hordeum vulgare subsp. Vulgare*, Lj – *Lemna japonica*, Os – *Oryza sativa Japonica*, Pta – *Populus tremula x Populus alba*, SI – *Solanum lycopersicum*, Sp - *S. polyrhiza*, St – *Solanum tuberosum*, Ta – *Triticum aestivum*, Wa – *Wolffia australiana*, Zm – *Zea mays*. Protein identifiers are listed in Table S1 (SUTs) and Table S2 (SWEETs). Phylogenies were calculated according to the nearest neighbor clustering method. A phylogenetic tree including the SWEET genes of *W. australiana*, *L. japonica* and *L. minor* is provided in Fig. S1.

Of the SWEETs, members of clades I, II, and III were identified in *S. polyrhiza* (Fig. 1B) and other duckweeds (Fig. S1). In clade I, there are homologues to Arabidopsis AtSWEET1 and AtSWEET2 (Fig. 1B) that were reported to transport glucose (Chen et al. 2010). In clade II, there are homologues to rice OsSWEET4 and rice OsSWEET6 (Fig. 1B) that were shown to transport sucrose (Sosso et al. 2015). Furthermore, in clade III, SpSWEET11 and SpSWEET12 are homologues to a group of SWEETs known to transport sucrose, including in active phloem loading, in Arabidopsis (Fig. 1B, Chen, 2014; Fatima et., 2022).

### Sucrose transporting SUTs and SWEETs are not expressed in the frond phloem

We tested the ability of selected SUT and SWEET transporters of S*. polyrhiza* to transport glucose and sucrose in a yeast experimental system (Fig. 2). The yeast strains either lacked the capacity to take up glucose or to utilize extracellular sucrose. The yeast growth assays indicated that SpSWEET1 facilitates glucose transport (Fig. 2A), while SpSUT2 and SpSWEET11 mediate sucrose transport (Fig. 2B). SpSWEET4 did not facilitate glucose or sucrose beyond background levels (Fig. 2A,B). The sucrose-transport ability of SpSUT2 was confirmed by application of the fluorescent sucrose analogue esculin. This was taken up into *SpSUT2*-expressing yeast cells in the expected pH- and substrate concentration-dependent manner, although at a much lower rate than by cells expressing *AtSUC2* (Fig. 2C,D).

**Fig. 2.**
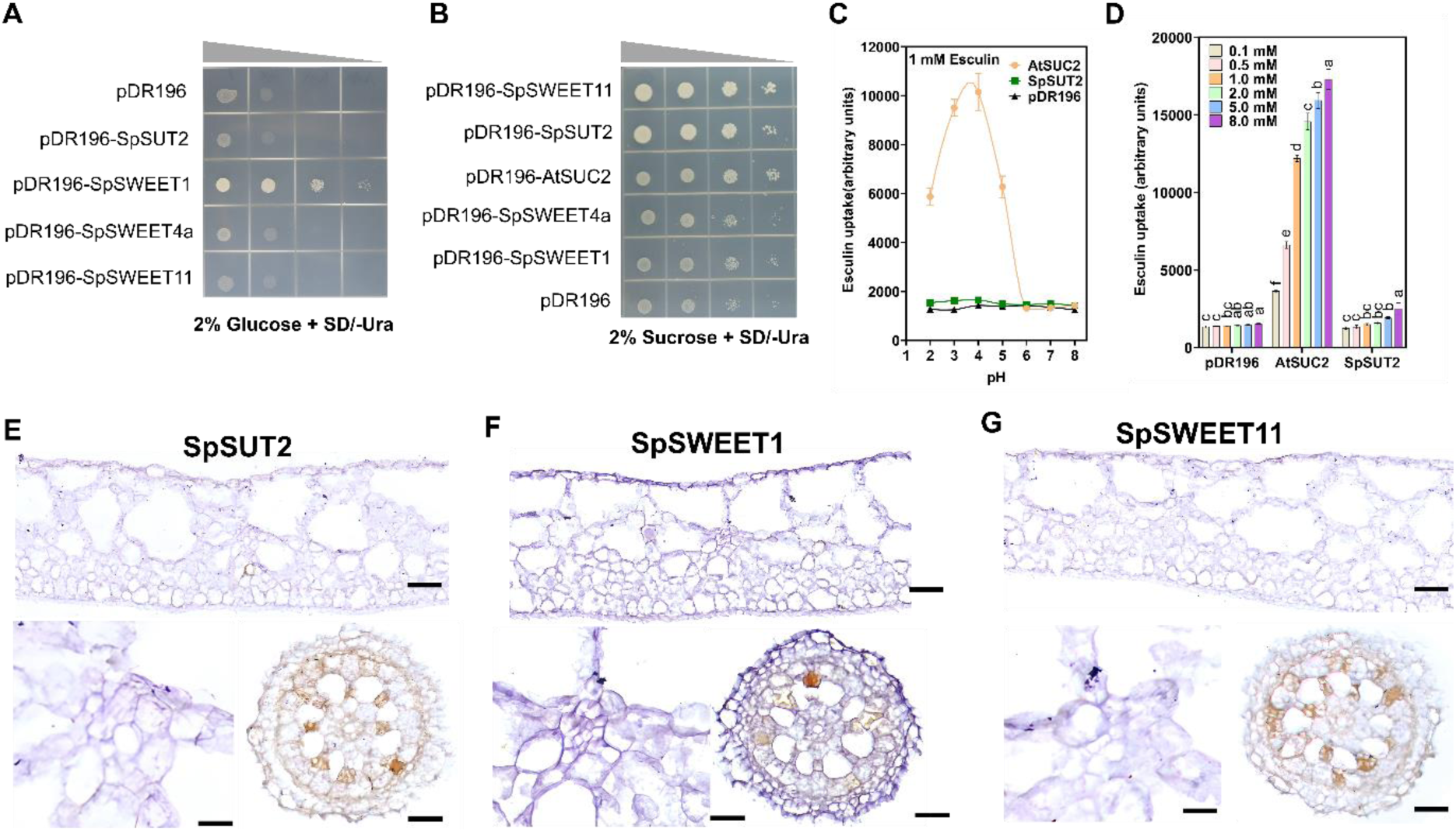
Analysis of transport function and *in situ* gene activity. (A) Growth of glucose uptake-deficient yeast (*EBY.VW4000*) transformed with the specified gene constructs or the empty vector (pDR196) in 2% glucose SD/-Ura medium. (B) Growth of sucrose uptake-deficient yeast (*SUSY7*/*ura3*) on 2% sucrose SD/-Ura medium. Arabidopsis AtSUC2 serves as positive control. Triangles in A and B indicate yeast concentration. The assays of A and B were each repeated three times and showed the same result each time. (C) pH dependence of esculin uptake in control and complemented yeast strains. (D) Concentration dependence of esculin uptake. Bars in (C) and (D) indicate standard deviation. Significance of difference tested by *t*-test (p < 0.05) is indicated by different letters. N = 3. (E-G) Tissue images after *in situ* hybridization with probes specific for SpSUT2 (E), SpSWEET1 (F), SpSWEET11 (G). Shown are a representative frond cross section (upper panel), a magnified view of a minor vein (lower left) and a cross section of the root (lower right). Brown staining indicates mRNA abundance. At least three different plants were analyzed for each gene, all showing the same staining pattern. Scale bars: 50 µm (frond and root sections), 10 µm (minor vein views).

To investigate if the sugar transport functions of SpSUT2, SpSWEET1 and SWEET11 relate to phloem loading, we checked their expression domain by *in situ* mRNA hybridization. mRNA of these transporters was not detected at the site where active phloem loading would take place, the minor veins (Fig. 2E-G). SpSUT2 activity was indicated in some frond mesophyll cells, as well as inner and middle cortical cells and epidermis cells in the root (Fig. 2E). SpSWEET1 activity was detected in root middle cortical cells (Fig. 2F). SpSWEET11 mRNA was detected in root inner and middle cortical cells (Fig. 2G). Cell type classification followed Kim (2007).

### SUT inhibition does not influence phloem exudate sucrose content

As in typical leaves, the hexoses fructose and glucose were more abundant than sucrose in the fronds of *S. polyrhiza* (Fig. 3A). In phloem exudate, sucrose was concentrated much higher than glucose and fructose (Fig. 3B). No raffinose or stachyose could be detected. A similar relative sugar distribution is found in all active apoplastic and passive symplastic phloem loaders, including all monocots analyzed so far (Zhang and Turgeon 2021). To differentiate between the two loading types, we applied the SUT inhibitor p-chloromercuribenzenesulfonic acid (pCMBS) to *S. polyrhiza* fronds. This did not affect the sucrose concentration in phloem exudate (Fig. 3B), further arguing against a role of SUTs in *S. polyrhiza* phloem loading.

**Fig. 3.**
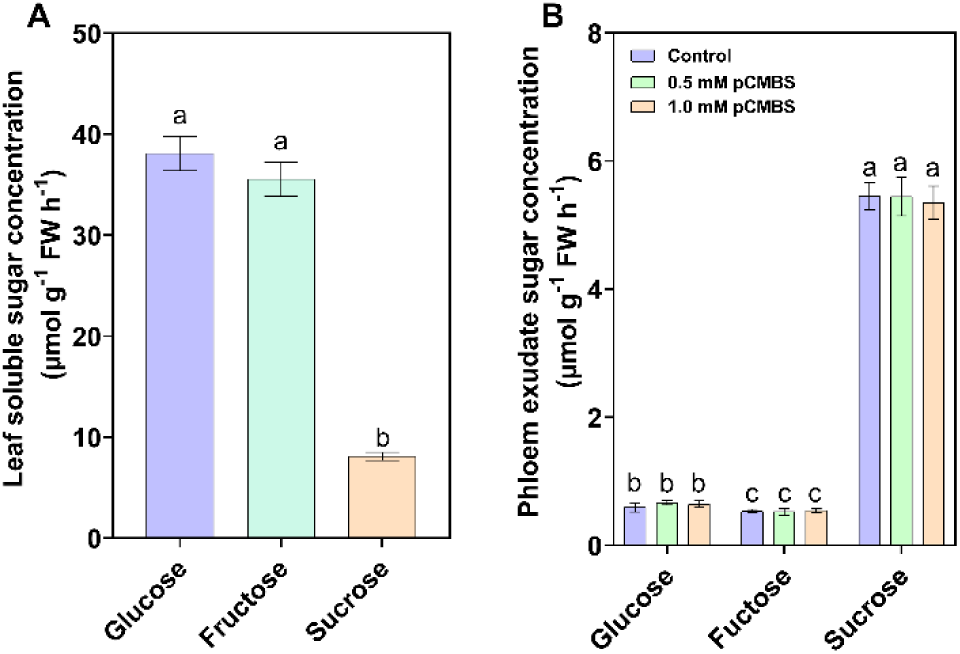
Soluble sugar concentration in frond extract (A) and phloem exudate (B). Phloem exudate sugar concentrations were measured without or x hours after application of the SUT inhibitor p-chloromercuribenzenesulfonic acid (pCMBS) at the specified concentration. Error bars indicate standard deviation. N = 3. Significance of difference tested by *t*-test (p < 0.05) is indicated by different letters.

### High plasmodesmata density along the phloem loading pathway

The relative density of plasmodesmata can serve as indicator of phloem loading type (Liesche et al. 2019). Active apoplastic loaders have much fewer plasmodesmata between companion cells and phloem parenchyma cells than at the other interfaces along the loading path. Active symplastic loaders, in contrast, have many times more plasmodesmata between companion cells and bundle sheath cells than at any other interface. In passive symplastic loaders plasmodesmata density is similar for all interfaces along the phloem loading pathway.

We counted plasmodesmata on electron micrographs of frond cross sections (Fig. 4). The combination of data from a high number of sections enabled the calculation of plasmodesmata density at the different cell-cell interfaces. The results were compared to average values for species with active apoplastic, active symplastic and passive symplastic loading derived from Liesche et al. (2019). With values between 50 and 60 plasmodesmata along 1 µm of vein, abundance in *S. polyrhiza* was relatively high at all interfaces (Fig. 4F). The relative density along the phloem loading pathway showed the limited variation typical for passive symplastic phloem loaders (Fig. 4G).

**Fig. 4.**
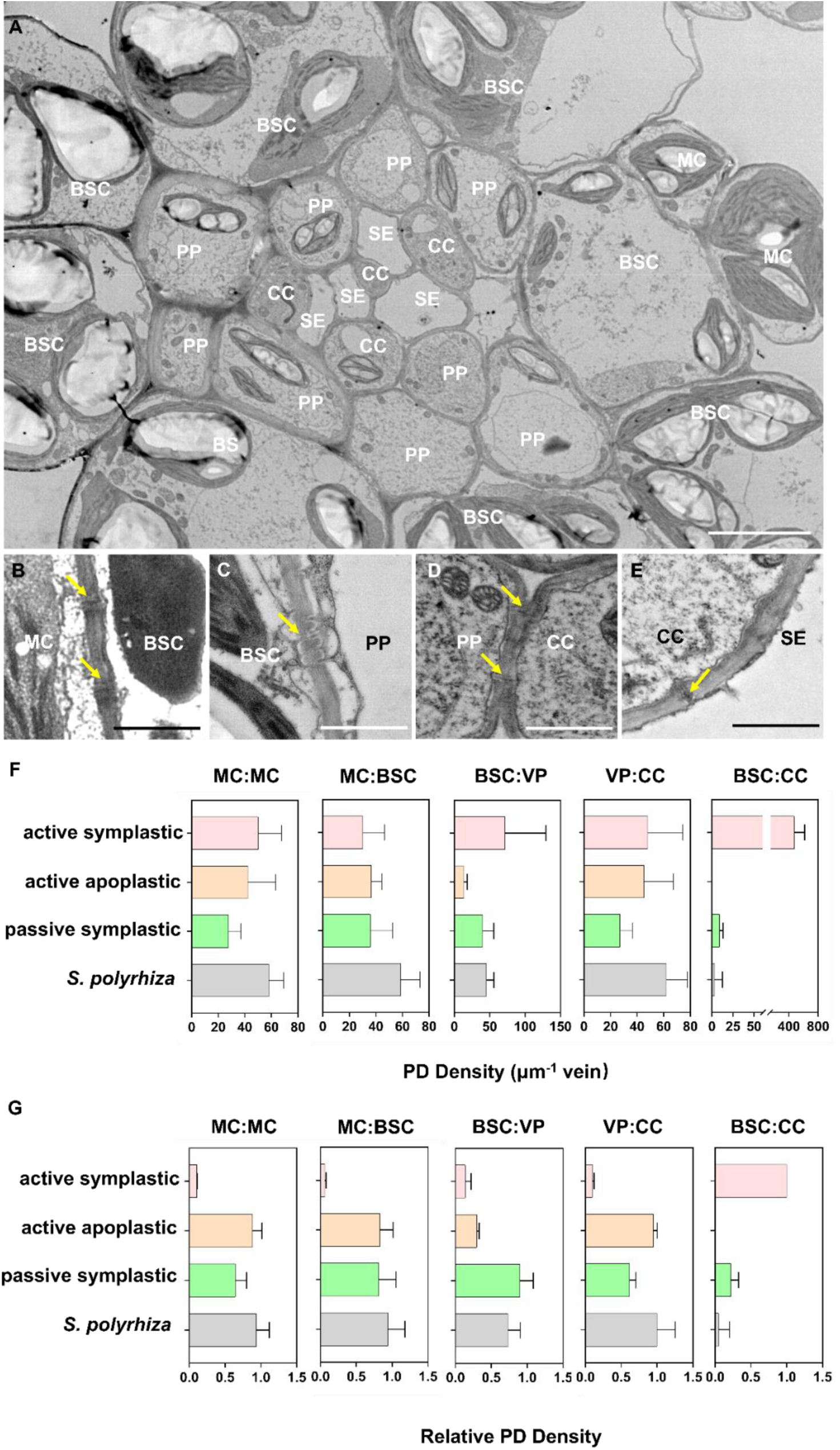
Plasmodesmata density analysis on electron micrographs of *S. polyrhiza* frond cross sections. (A) overview image of a minor vein featuring bundle sheath cells (BSC), mesophyll cells (MC), phloem parenchyma cells (PP), companion cells (CC) and sieve elements (SE). (B-E) Magnifications of plasmodesmata (yellow arrows) at the different interfaces along the phloem loading pathway: MC to BSC (B), BSC to PP (C), PP to CC (D), CC to SE (E). (F) Plasmodesmata density calculated as the number of PD per interface per 1 µm of minor vein. (G) Relative plasmodesmata density, the number of plasmodesmata at each interface normalized to the highest value along the phloem-loading pathway. Error bars indicate standard deviation. Plasmodesmata were counted on 15 minor veins from 12 different plants.

### Hormone-induced modification of plasmodesmata co-occurs with changes in phloem sugar content

The above results all indicate passive phloem loading through plasmodesmata in *S. polyrhiza*. Accordingly, the permeability of plasmodesmata should influence the loading rate. To test this hypothesis, we looked for substances that would influence the permeability of plasmodesmata in the fronds. We chose several hormones that have been previously implicated in plasmodesmata regulation in other plant species (Han and Kim 2016; Sankoh and Birch-Smith 2021). In fluorescence redistribution after photobleaching (FRAP) experiments on living frond epidermis cells, their influence on plasmodesmata-mediated interface permeability was tested. The mobile fraction of the fluorescent tracer was relatively low at about 20 % across experiments (Fig. 5). Significant differences were recorded for the rate of mobility, indicated by the half-time of recovery. In control plants, the mean half-time was about 2.2 s (Fig. 5). While it was slightly reduced after GA treatment (2.08 s), half-times were increased after ABA (2.53 s), NAA (2.5 s) and SA (2.32s) treatments (Fig. 5). These longer half-times are indicative of reduced transport capacity, potentially through reduced plasmodesmata permeability.

**Fig. 5.**
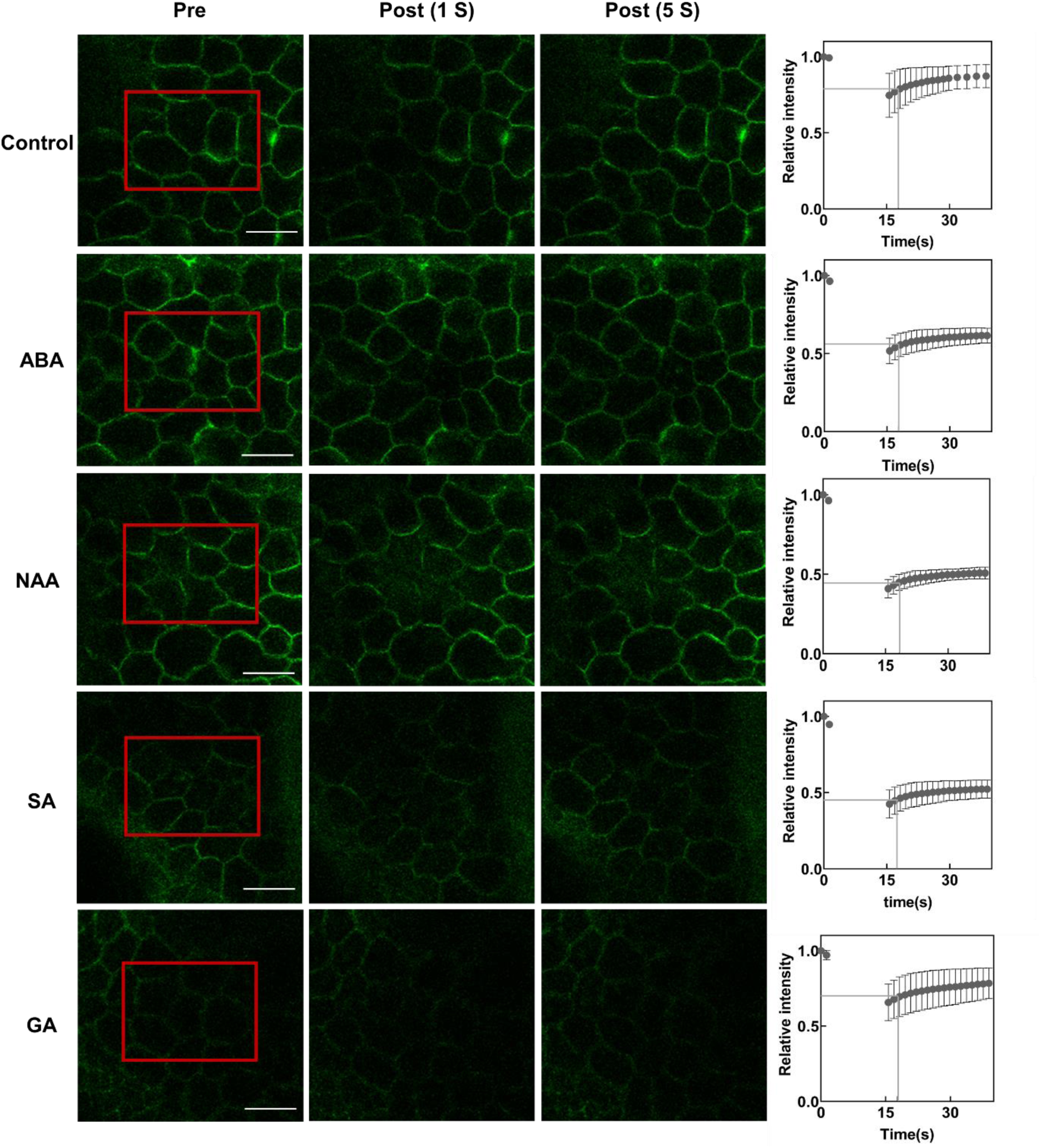
Fluorescence redistribution after photobleaching (FRAP) experiments indicate plasmodesmata-mediated interface permeability in *S. polyrhiza* frond epidermis cells. The fluorescence images represent the FRAP experiments conducted 12 hours after moving plants to control or hormone-containing medium. From left to right, a frond area is shown before bleaching (Pre), 1 s after the 15 s-long bleaching (Post 1 s) and 5 s after the bleaching (Post 5 s). ABA – abscisic acid, NAA – 1-naphthaleneacetic acid, SA – salicylic acid, GA – gibberellic acid. Scale bars: 50 µm. The diagrams on the right show the relative fluorescence intensity with bleaching happening between 2 s and 17 s timepoints. Data points indicate means of 3 experiments. Error bars indicate standard deviation. Grey lines indicate half-life.

The conclusions drawn from FRAP experiments were partly corroborated by aniline blue staining assays (Fig. 6). Aniline blue fluorescence serves as indicator of callose levels at plasmodesmata (Zavaliev and Epel 2015). More plasmodesmata callose is associated with lower permeability (Amsbury et al. 2017). The treatment with ABA and NAA, which decreased fluorescence redistribution in the FRAP experiment the most, showed the strongest increase in aniline blue fluorescence (Fig. 6A-F). GA treatment led to lower aniline blue fluorescence levels (Fig. 6E,F). SA treatment did not significantly change aniline blue fluorescence compared to the control (Fig. 6F).

**Fig. 6.**
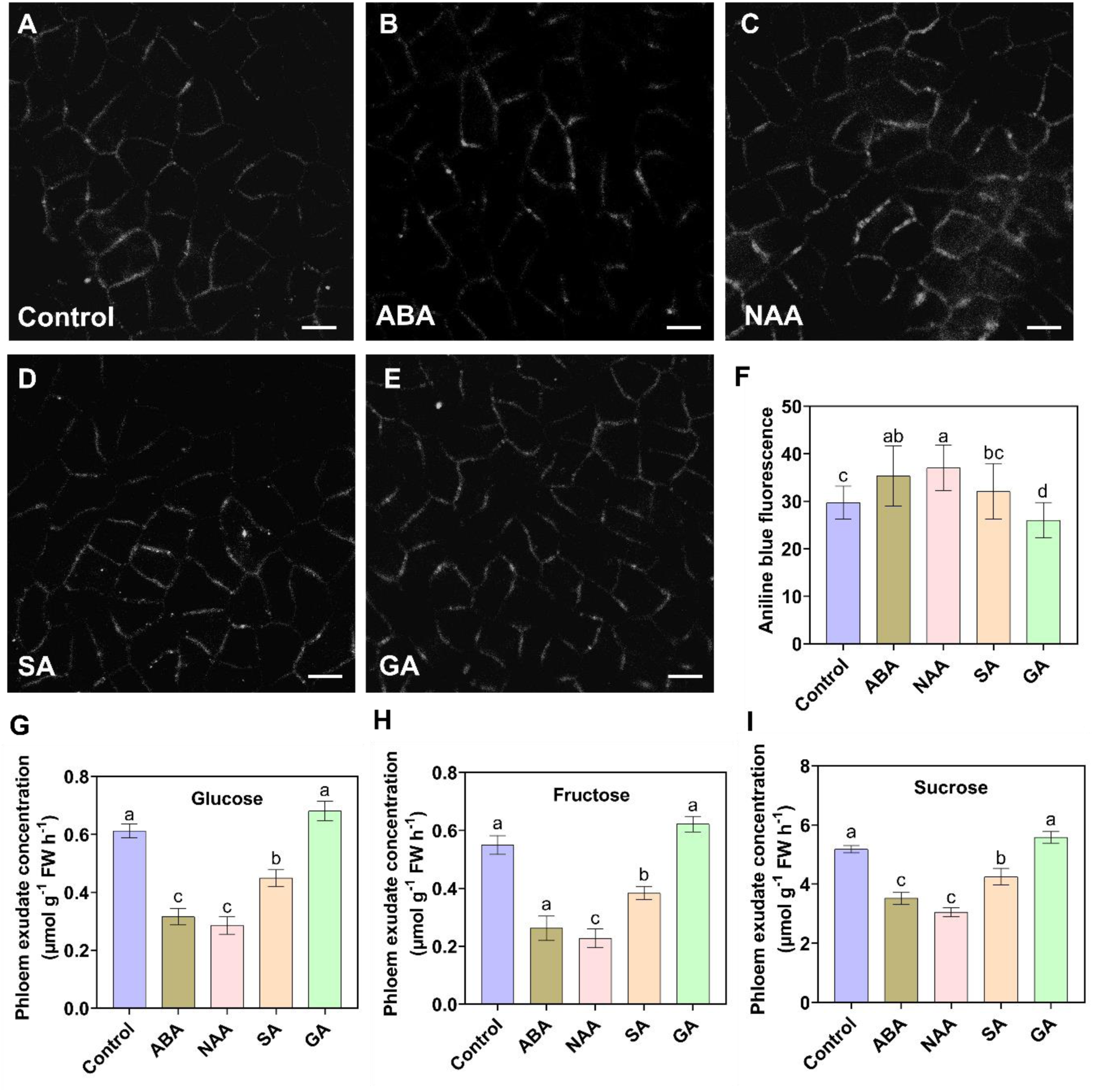
Hormone treatment effect on aniline blue staining and phloem exudate sugar concentrations. (A-E) Fluorescence images of aniline blue-stained frond epidermis cells indicating plasmodesmata callose levels. Scale bars 10 µm. (F) Quantification of aniline blue fluorescence. (G-I) Phloem exudate concentrations of glucose (G), fructose (H) and sucrose (I). ABA – abscisic acid, NAA – 1-naphthaleneacetic acid, SA – salicylic acid, GA – gibberellic acid. Error bars indicate standard deviation. N = 30 (F), 3 (G-I). Significance of difference tested by *t*-test (p < 0.05) is indicated by different letters.

The effect of hormone treatment on callose levels was further evaluated by measurement of mRNA levels of callose synthases and β-glucanases. β-glucanases, also called glucan endo-1,3-beta-glucosidase, are responsible for callose breakdown. mRNA sequencing of frond tissue after hormone treatment showed callose synthase *SpGALS2* strongly upregulated, while the other callose synthase in the *S. polyrhiza* genome, *SpGALS1*, was strongly downregulated after treatment with ABA, NAA and SA (Table 1). Three of the four β-glucanases were down-regulated (Table 1). GA treatment, in contrast, did not affect callose synthase mRNA levels and only led to down-regulation of one β-glucanase (Table 1).

**Table 1.** Change of mRNA levels of selected genes in *S. polyrhiza* plants after hormone treatment. ABA – abscisic acid, NAA – 1-naphthaleneacetic acid, SA – salicylic acid, GA – gibberellic acid. The p-value indicates statistical significance of the change for a single gene. The q-value is an adjusted p-value that considers the false discovery rate. A q-value < 0.05 is considered statistically significant, at which 5% of the tests with p < 0.05 are assumed to be false positives.

| Gene | fold change | log2-fold change | p-value | q-value | Significant regulation |
| --- | --- | --- | --- | --- | --- |
| **ABA_vs_control** |  |  |  |  |  |
| SpSUT2<br>(Spipo11G0051900) | 1.10569 | 0.14495 | 0.09402 | 0.1185 |  |
| SpSUT4<br>(Spipo2G0106800) | 3.10595 | 1.63503 | 0.00045 | 0.00076 | Up |
| SpSWEET1<br>(Spipo4G0094000) | 0.91660 | -0.12563 | 0.15283 | 0.18532 |  |
| SpSWEET2a<br>(Spipo25G0000900) | 0.83525 | -0.25971 | 1.76633E-06 | 3.57182E-06 | Down |
| SpSWEET4a<br>(Spipo26G0025500) | 0.02032 | -5.62037 | 2.66565E-11 | 7.14622E-11 | Down |
| SpSWEET11<br>(Spipo4G0059100) | 0.02537 | -5.30018 | 4.1984E-06 | 8.26791E-06 | Down |
| SpSWEET12<br>(Spipo0G0112200) | 1.75857 | 0.81440 | 0.52711 | 0.57133 |  |
| SpGALS2<br>(Spipo5G0045900) | 2.69429 | 1.42990 | 7.46243E-35 | 4.2409E-34 | Up |
| SpGALS1<br>(Spipo12G0041000) | 0.24659 | -2.01979 | 4.5107E-101 | 6.5869E-100 | Down |
| Spipo3G0064700<br>(β-glucanase) | 0.0029 | -8.38530 | 3.69702E-12 | 1.03016E-11 | Down |
| Spipo3G0064800<br>(β-glucanase) | 0.38665 | -1.37087 | 8.1146E-162 | 1.9294E-160 | Down |
| Spipo6G0010200<br>(β-glucanase) | 0.25366 | -1.97900 | 0.1287 | 0.15830 |  |
| Spipo6G0010300<br>(β-glucanase) | 0.33999 | -1.55641 | 4.3309E-195 | 1.3072E-193 | Down |
| **NAA_vs_control** |  |  |  |  |  |
| SpSUT2<br>(Spipo11G0051900) | 1.60640 | 0.6838 | 4.4724E-17 | 1.55143E-16 | Up |
| SpSUT4<br>(Spipo2G0106800) | 0.84880 | -0.23648 | 0.71459 | 0.75593 |  |
| SpSWEET1<br>(Spipo4G0094000) | 0.61190 | -0.70861 | 4.26579E-14 | 1.32824E-13 | Down |
| SpSWEET2a<br>(Spipo25G0000900) | 0.60614 | -0.72226 | 6.80685E-35 | 3.72061E-34 | Down |
| SpSWEET4a<br>(Spipo26G0025500) | 0.29555 | -1.75849 | 1.61846E-11 | 4.55601E-11 | Down |
| SpSWEET11<br>(Spipo4G0059100) | 0.07336 | -3.76885 | 5.20468E-06 | 1.07527E-05 | Down |
| SpSWEET12<br>(Spipo0G0112200) | 1.96081 | 0.97145 | 0.46919 | 0.52562 |  |
| SpGALS2<br>(Spipo5G0045900) | 2.8038 | 1.51120 |  |  | Up |
| SpGALS1<br>(Spipo12G0041000) | 0.35299 | -1.50226 | 4.10121E-65 | 3.80138E-64 | Down |
| Spipo3G0064700<br>(β-glucanase) | 0.27436 | -1.8658 | 8.84463E-10 | 2.28339E-09 | Down |
| Spipo3G0064800<br>(β-glucanase) | 0.29732 | -1.74987 | 2.1345E-221 | 6.8376E-220 | Down |
| Spipo6G0010200<br>(β-glucanase) | 0.67354 | -0.57015 | 0.60928 | 0.65992 | |
| Spipo6G0010300<br>(β-glucanase) | 0.2024 | -2.30427 | 0 | 0 | Down |
| **SA_vs_control** |  |  |  |  |  |
| SpSUT2 (<br>Spipo11G0051900) | 1.03110 | 0.04418 | 0.60449 | 0.65932 |  |
| SpSUT4<br>(Spipo2G0106800) | 1.51634 | 0.6005 | 0.25508 | 0.31108 |  |
| SpSWEET1<br>(Spipo4G0094000) | 0.46412 | -1.10742 | 2.24404E-32 | 1.65466E-31 | Down |
| SpSWEET2a<br>(Spipo25G0000900) | 1.21396 | 0.27972 | 1.18931E-07 | 3.10248E-07 | Up |
| SpSWEET4a<br>(Spipo26G0025500) | 0.64796 | -0.6260 | 0.0044 | 0.00758 | Down |
| SpSWEET11<br>(Spipo4G0059100) | 0.03615 | -4.78970 | 9.74922E-07 | 2.38993E-06 | Down |
| SpSWEET12<br>(Spipo0G0112200) | 2.24333 | 1.16564 | 0.33709 | 0.39855 |  |
| SpGALS2<br>(Spipo5G0045900) | 2.4025 | 1.26454 | 4.42629E-27 | 2.79716E-26 | Up |
| SpGALS1<br>(Spipo12G0041000) | 0.35408 | -1.4978 | 9.18322E-82 | 1.62369E-80 | Down |
| Spipo3G0064700<br>(β-glucanase) | 0.04496 | -4.47491 | 2.74478E-24 | 1.59144E-23 | Down |
| Spipo3G0064800<br>(β-glucanase) | 0.10215 | -3.29115 | 0 | 0 | Down |
| Spipo6G0010200<br>(β-glucanase) | 0.78318 | -0.35257 | 0.71885 | 0.7619 | |
| Spipo6G0010300<br>(β-glucanase) | 0.50736 | -0.97889 | 6.16727E-87 | 1.15487E-85 | Down |
| **GA_vs_control** |  |  |  |  |  |
| SpSUT2<br>(Spipo11G0051900) | 1.07272 | 0.10127 | 0.23833 | 0.44137 |  |
| SpSUT4<br>(Spipo2G0106800) | 0.8811 | -0.18257 | 0.76378 | 0.87483 |  |
| SpSWEET1<br>(Spipo4G0094000) | 1.128236139 | 0.174069055 | 0.027205378 | 0.095126443 |  |
| SpSWEET2a<br>(Spipo25G0000900) | 0.951938205 | -0.071060171 | 0.199313617 | 0.391979518 |  |
| SpSWEET4a<br>(Spipo26G0025500) | 0.797944418 | -0.325639837 | 0.11209685 | 0.264782127 |  |
| SpSWEET11<br>(Spipo4G0059100) | 1.121943953 | 0.166000607 | 0.700842039 | 0.83641036 |  |
| SpSWEET12<br>(Spipo0G0112200) | 0.844050161 | -0.244599355 | 0.934050314 | 0.969025286 |  |
| SpGALS2 | 0.834129908 | -0.261656007 | 0.07148517 | 0.193961626 |  |
| (Spipo5G0045900) |  |  |  |  |  |
| SpGALS1 | 0.927099131 | -0.109204486 | 0.111429348 | 0.263988005 |  |
| (Spipo12G0041000) |  |  |  |  |  |
| Spipo3G0064700 | 1.13695411 | 0.185174025 | 0.333186804 | 0.545378447 |  |
| (β-glucanase) | | | | | |
| Spipo3G0064800 | 0.815009334 | -0.295111513 | 3.93944E-10 | 9.5673E-09 | Down |
| (β-glucanase) | | | | | |
| Spipo6G0010200 | 0.580342317 | -0.785023964 | 0.462671272 | 0.666119632 |  |
| (β-glucanase) | | | | | |
| Spipo6G0010300 | 1.007520231 | 0.010808808 | 0.825571639 | 0.911232865 |  |
| (β-glucanase) | | | | | |

Considering the clear changes of plasmodesmata permeability, especially after treatment with ABA and NAA, we tested if these affect phloem loading. Indeed, phloem exudate sugar levels reacted to most hormone treatments. Hexoses and sucrose were strongly decreased by ABA and NAA treatment, while SA had a smaller effect (Fig. 6G-I). GA did not affect phloem exudate sugar levels (Fig. 6G-I). Thereby, the hormone effects on phloem exudate sugar levels were similar to the effects on frond plasmodesmata permeability. SUT and SWEET mRNA levels showed no consistent reactions to the hormone treatments (Table 1).

## Discussion

Duckweeds are aquatic plants that are generally recognized as the smallest angiosperms (Ziegler et al. 2014). Accordingly, their phloem could serve different purposes than in the duckweed’s well-studied monocot relatives like maize or rice. Especially, since a recent experiment showed a limited influence of root removal on frond growth (Ware et al. 2023). However, the presence of phloem in fronds and roots with similar anatomy to grasses (Fig. 4; Kim 2007) indicates that it shares the core function of long-distance sugar transport. That sucrose is transported in the phloem was confirmed here for *S. polyrhiza* by phloem exudate analysis (Fig. 3). It also showed sucrose to be the primary sugar, like in the grasses (Zhang and Turgeon 2018).

We can speculate that, besides supporting root growth and development, fast sucrose transport in the phloem might also be needed for sugar secretion. The root microbiome has immense influence on the growth of duckweeds (Acosta et al. 2021) and controlled sugar secretion could be employed to maintain a beneficial microbiome (Song et al. 2022; Loo et al. 2024). Moreover, duckweeds are able to accumulate high amounts of starch in root cells (Jones et al. 2021). Fast sucrose transport from frond to root could prevent feedback inhibition of photosynthesis, by providing a path to carbon storage in root cells. Therefore, understanding how sucrose is transported in the duckweed phloem could facilitate further improvements for their biotechnological use.

The results of our experiments all indicate that sucrose is loaded into the *S. polyrhiza* phloem in a passive symplastic manner. Their genome does not contain homologues to the SUTs associated with active apoplastic phloem loading in other monocots (Fig. 1). The other SUTs and the sucrose transporting SWEETs were not expressed in phloem minor veins, but mainly in root cells (Fig. 2). In the duckweed *Landoltia punctata* the *SUT2* promoter was active primarily in the guard cells (Liu et al. 2022), which we did not detect for *S. polyrhiza*. The primary function of the duckweed SUTs and SWEETs could lie in short-distance carbon allocation (Denyer et al. 2024). Further localization of expression at different developmental stages and under different growth conditions, as well as the analysis of plants with artificially modified gene activity, could provide further insight into their role. Active apoplastic loading in *S. polyrhiza* is further excluded based on the missing influence of the SUT-inhibitor pCMBS on the phloem exudate sucrose content (Fig. 3). Application of pCMBS to maize leaves caused a decrease in phloem loading (Thompson and Dale, 1981; Thorpe and Minchin, 1988). Active symplastic phloem loading can be ruled out based on the absence of sugar oligomers in the *S. polyrhiza* phloem exudate and the absence of highly abundant Y-shaped plasmodesmata between bundle-sheath and companion cells (Fig. 4). A clear indication for passive symplastic phloem loading in *S. polyrhiza* is the high, and evenly distributed, symplastic capacity at all cell interfaces between mesophyll cells and the companion cell-sieve element complex (Fig. 4).

Active phloem loading enables plants to keep low sugar levels in leaves, thereby facilitating efficient whole-plant carbon use (Turgeon 2010). The passive loading observed in many tree species, and the associated higher leaf sugar levels, reflect the optimization for long-term survival instead of maximal growth rates (Turgeon 2010). Duckweeds are considered the fastest growing plants (Acosta et al. 2021). However, their growth potential has only a minor dependence on phloem-mediated sugar transport, because growth mainly happens in the fronds. Accordingly, passive phloem loading is unlikely to have the same limiting effect on whole-plant growth potential observed in land plants. Investigating how much sucrose is transported in the phloem, and under what conditions, will be essential for understanding how sucrose phloem transport is integrated with whole-plant carbon usage in duckweeds.

In trees, the sucrose transport rate in the phloem is often seen as a simple function of supply and demand in the different parts of the plant (Sala et al. 2012; Guillemot et al. 2015). However, there are several examples of short-term regulation, in which leaf sucrose export rates are partly coupled from leaf sucrose production and sink demand (Liesche 2017). This could happen via dynamic regulation of plasmodesmata permeability (Ayyoub et al. 2024). Mathematical modeling showed that moderate changes in PD permeability would have a meaningful effect on the diffusion of sucrose into the phloem of passive loaders (Liesche 2017). Moreover, an observed increase in sucrose transport in tobacco leaves in response to JA application has been associated with increased PD permeability along the pre-phloem pathway (Han and Kim 2016). Our results, that reduced PD permeability coincides with reduced phloem exudate sucrose concentration (Fig. 5, Fig. 6) indicates that the dynamic regulation of PD could indeed influence the rate of sucrose loading in passive symplastic phloem loaders. However, there are a few issues to be considered regarding our results. Firstly, the hormones that were applied to change plasmodesmata permeability can potentially influence many other processes as well, e.g. photosynthesis or carbon management (Liu et al. 2019), meaning that the observed differences in phloem loading could be only partly related, or unrelated, to plasmodesmata permeability. Secondly, we measured plasmodesmata permeability only in epidermis cells because of experimental limitations. Different treatment effects of plasmodesmata in different cell types is possible (Sager et al. 2020). The experiments should be followed up by callose quantification in whole fronds at high spatial and temporal resolution, for example utilizing the ClearSee clearing technique (Kalmbach et al. 2023). Moreover, introduction of the inducible CalS3 system would enable the analysis of PD function partly independent from hormone function (Yan 2022). The reward will be a better understanding of passive phloem loading dynamics, not only in duckweeds, but also the many trees and shrubs that use this loading type.

## Acknowledgements

We sincerely appreciate the donation of the duckweed experimental materials by Associate Professor Liu Yu of Qingdao Agricultural University and Associate Researcher Xu Qiyu of Qingdao Institute of Bioenergy and Bioprocess Technology, Chinese Academy of Sciences. We thank the Northwest A&F University North Campus Life Science Core Services facility for supporting our imaging efforts.

---

**Fig. S1:**
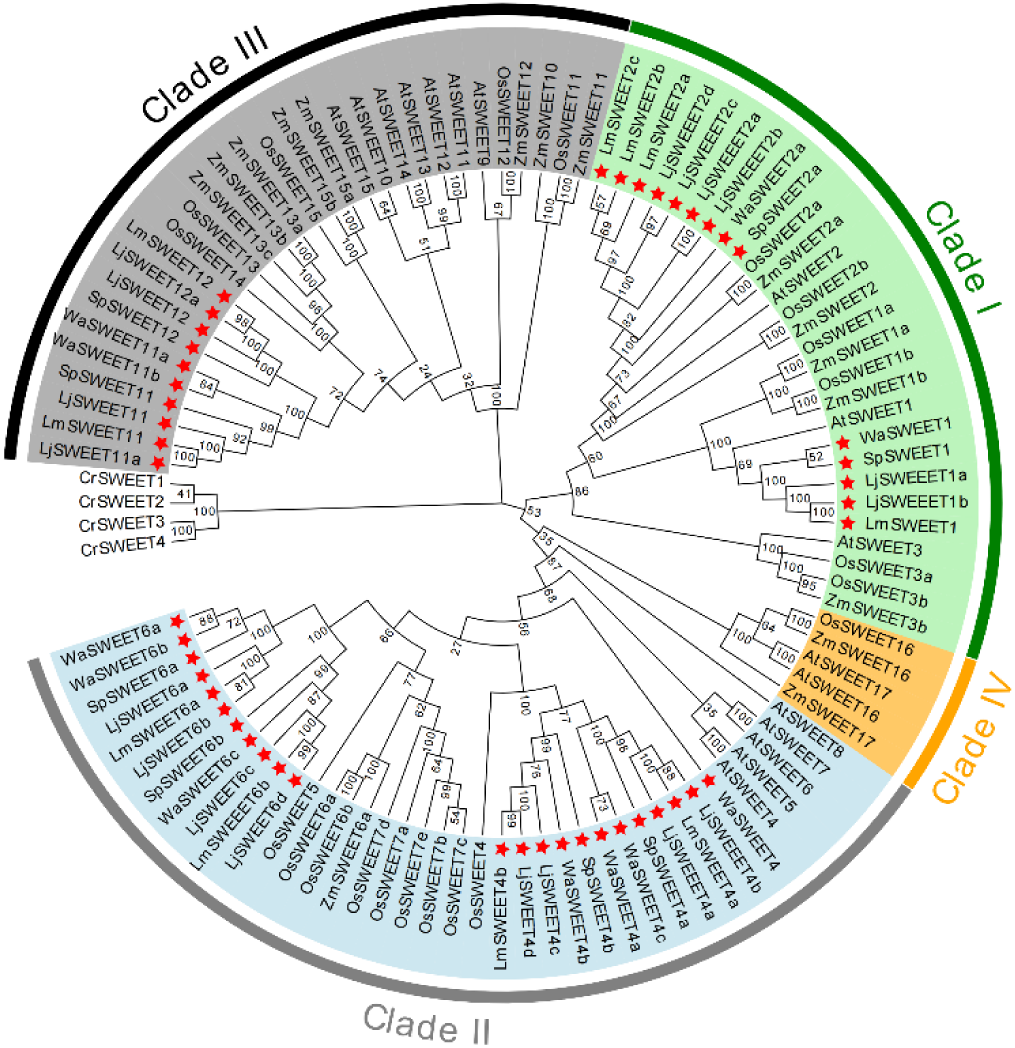
Phylogenetic analysis of SWEET gene family based on protein sequences. Duckweed genes are marked by stars. Species abbreviations: At – *Arabidopsis thaliana*, Cr –*Chlamydomonas reinhardtii*, Gm – *Glycine max (soybean)*, Hv – *Hordeum vulgare subsp. Vulgare*, Lj – *Lemna japonica*, Os – *Oryza sativa Japonica*, Pta – *Populus tremula x Populus alba*, SI – *Solanum lycopersicum*, Sp - *S. polyrhiza*, St – *Solanum tuberosum*, Ta – *Triticum aestivum*, Wa – *Wolffia australiana*, Zm – *Zea mays*. Protein identifiers are listed in Table S2.

**Supplementary Table S1.** Proteins used in the phylogenetic tree of SUTs

| Protein name | Species | Protein Accession number |
| --- | --- | --- |
| AcNRRL1 | *Aspergillus clavatus* | XP_001270868 |
| AtSUC1 | *Arabidopsis thaliana* | Q39232 |
| AtSUC2 | *Arabidopsis thaliana* | Q39231 |
| AtSUC3 | *Arabidopsis thaliana* | O80605 |
| AtSUC4 | *Arabidopsis thaliana* | Q9FE59 |
| AtSUC5 | *Arabidopsis thaliana* | Q9C8X2 |
| AtSUC6 | *Arabidopsis thaliana* | Q6A329 |
| AtSUC7 | *Arabidopsis thaliana* | Q67YF8 |
| AtSUC8 | *Arabidopsis thaliana* | Q9ZVK6 |
| AtSUC9 | *Arabidopsis thaliana* | Q9FG00 |
| OsSUT1 | *Oryza sativa Japonica Group* | Q10R54 |
| OsSUT2 | *Oryza sativa Japonica Group* | Q0ILJ3 |
| OsSUT3 | *Oryza sativa Japonica Group* | Q948L0 |
| OsSUT4 | *Oryza sativa Japonica Group* | Q6YK44 |
| OsSUT5 | *Oryza sativa Japonica Group* | Q69JW3 |
| ZmSUT1 | *Zea mays* | NP_001292720 |
| ZmSUT2 | *Zea mays* | NP_001137486 |
| ZmSUT4 | *Zea mays* | AAT51689 |
| PtaSUT1 | *Populus tremula x Populus alba* | ADW94615 |
| PtaSUT3 | *Populus tremula x Populus alba* | ADW94616 |
| PtaSUT4 | *Populus tremula x Populus alba* | ADW94617 |
| PtaSUT5 | *Populus tremula x Populus alba* | ADW94618 |
| PtaSUT6 | *Populus tremula x Populus alba* | ADW94619 |
| GmSUC1 | *Glycine max (soybean)* | Q7XA53 |
| HvSUT1 | *Hordeum vulgare subsp. vulgare* | Q4A1H2 |
| HvSUT4 | *Hordeum vulgare subsp. vulgare* | Q9M423 |
| SISUT1 | *Solanum lycopersicum* | BAO96213 |
| SISUT2 | *Solanum lycopersicum* | AAG12987 |
| SISUT4 | *Solanum lycopersicum* | AAG09270 |
| TaSUT1 | *Triticum aestivum* | AAM13408 |
| StSUT1 | *Solanum tuberosum* | Q43653 |
| StSUT2 | *Solanum tuberosum* | AAP43631 |
| SpSUT2 | *Spirodela polyrhiza* | Spipo11G0051900 |
| SpSUT4 | *Spirodela polyrhiza* | Spipo2G0106800 |
| LmSUT2 | *Lemna minor 7210* | Lm7210a004g010820_T003 |
| LmSUT4 | *Lemna minor 7210* | Lm7210a005g002070_T001 |
| LjSUT2 | *Lemna japonica 9421* | Lj9421a04Tg010320_T001 |
| LjSUT4 | *Lemna japonica 9421* | Lj9421a05Mg002090_T002 |
| WaSUT2 | *Wolffia australiana 8730* | Wa8730c014g001690_T001 |
| WaSUT4 | *Wolffia australiana 8730* | Wa8730c003g001890_T001 |

**Supplementary Table S2.**
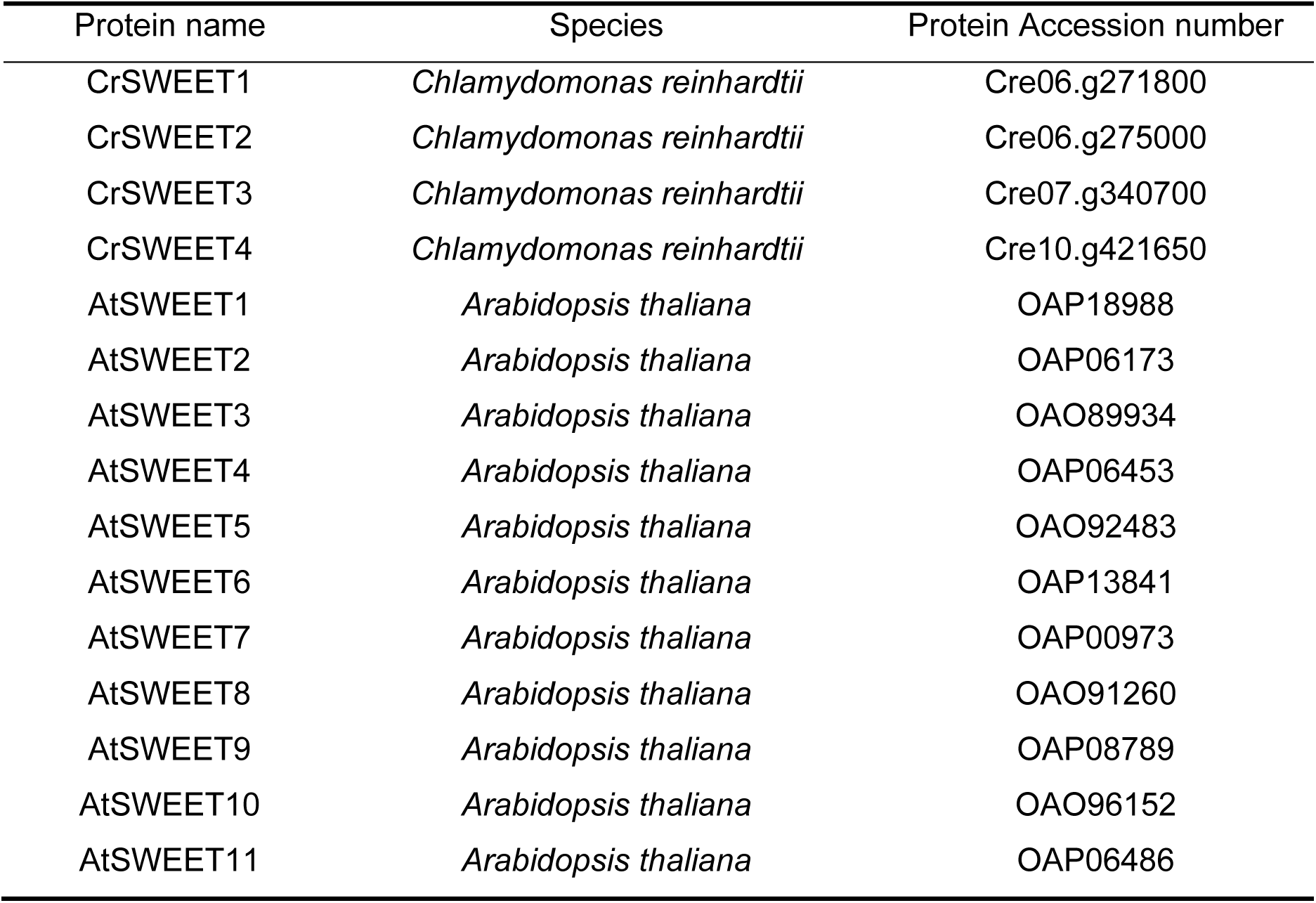
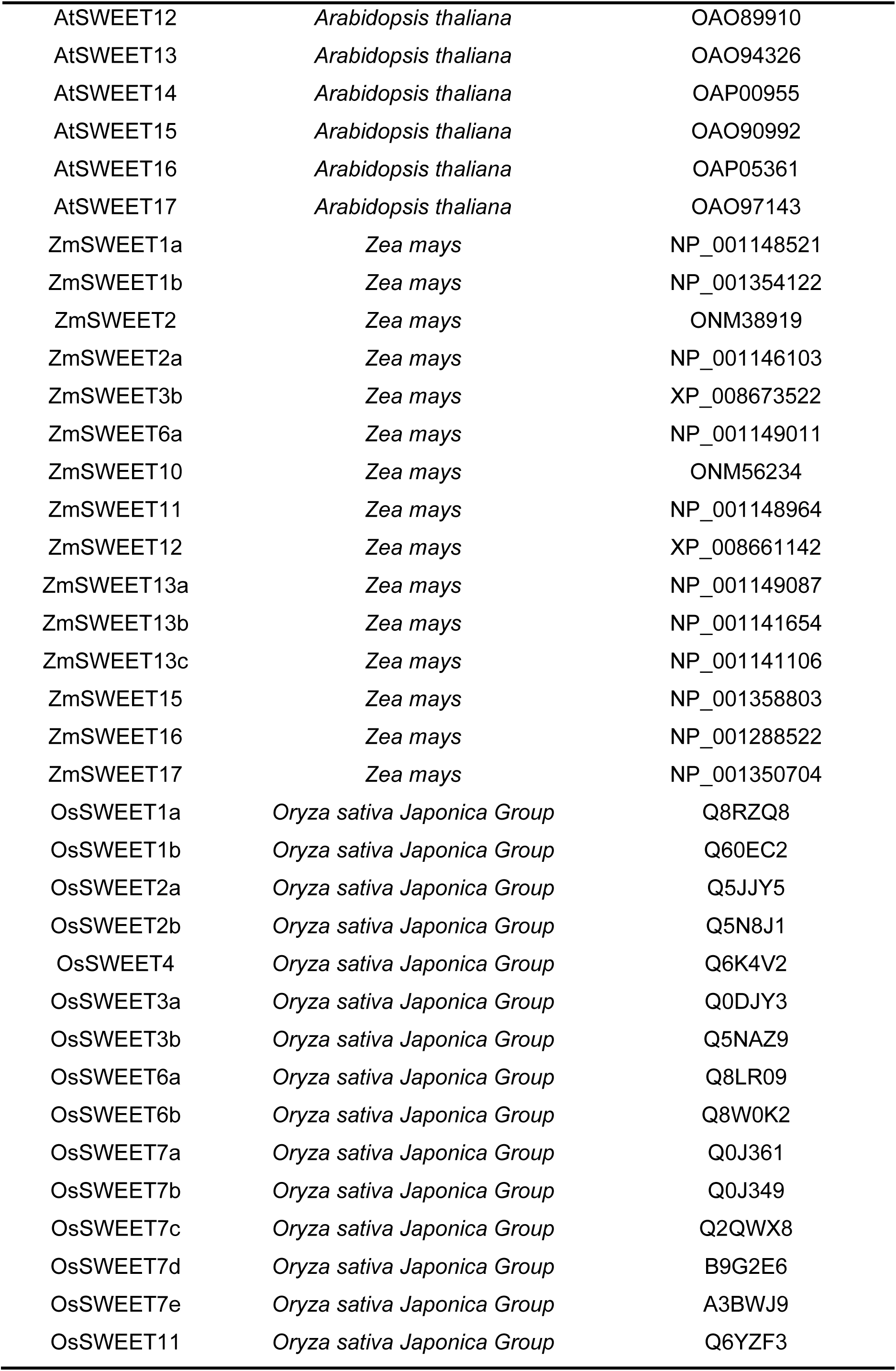
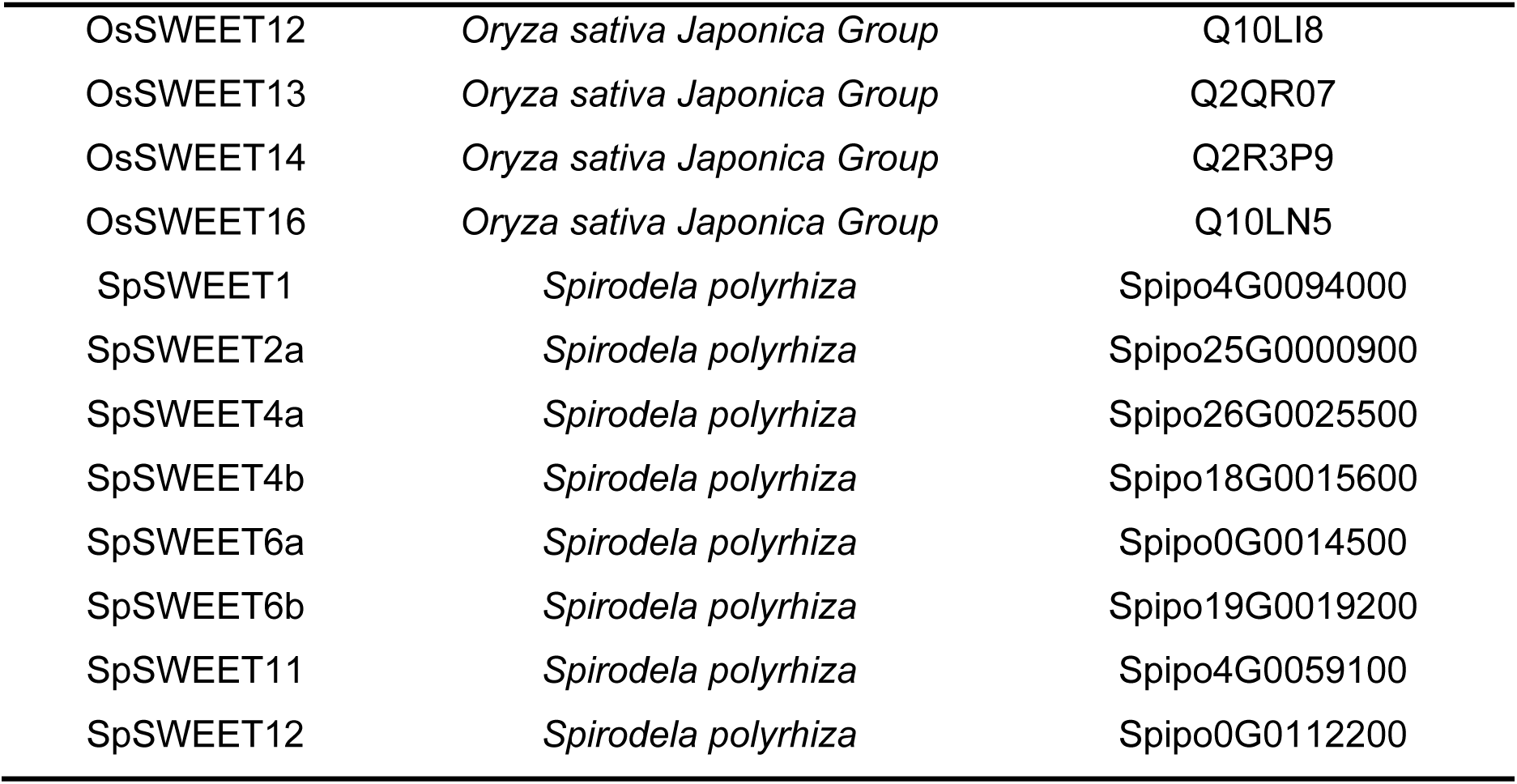
Proteins used in the phylogenetic tree of SWEETs

**Supplemental Table S3:**
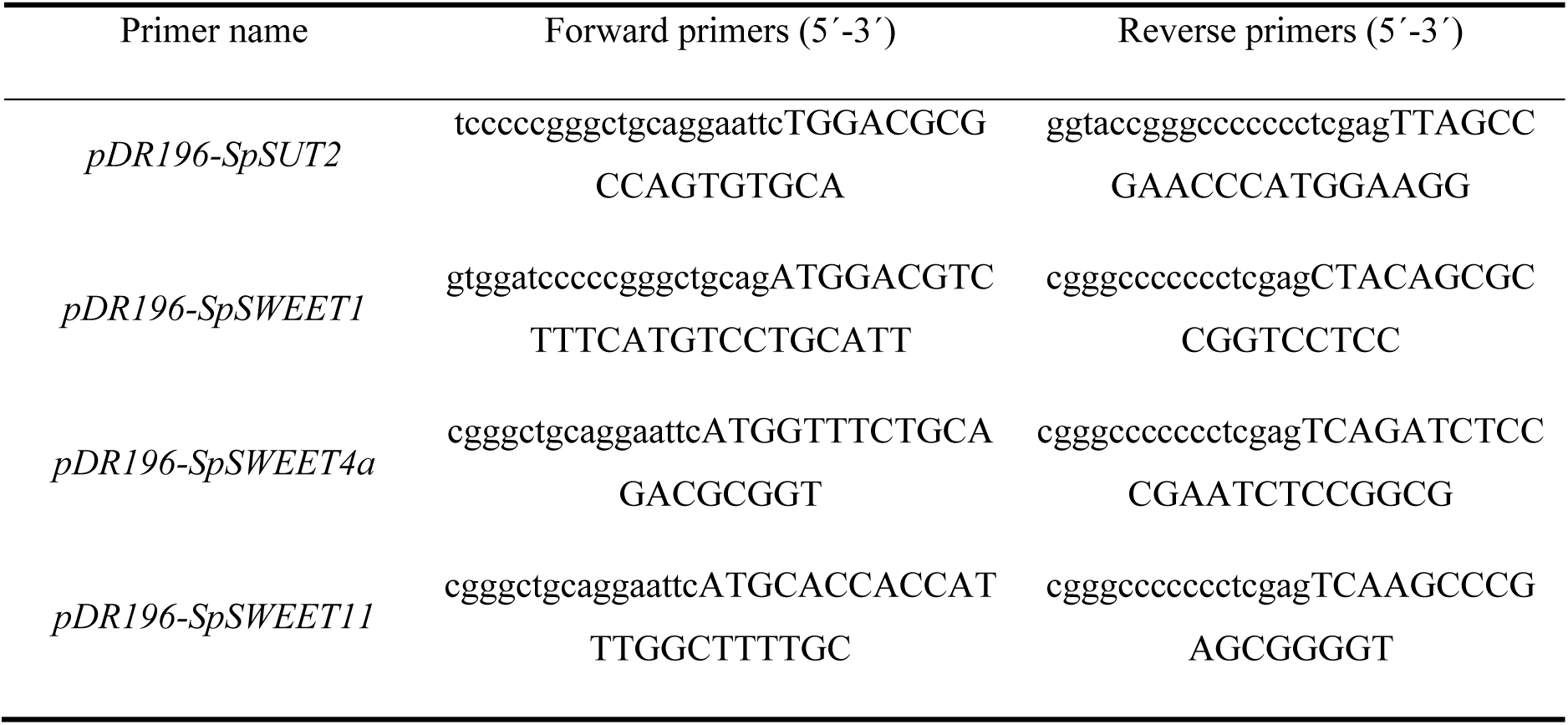
Primers for cloning of transporter genes into the pDR196 yeast expression vector

**Supplemental Table S4:**
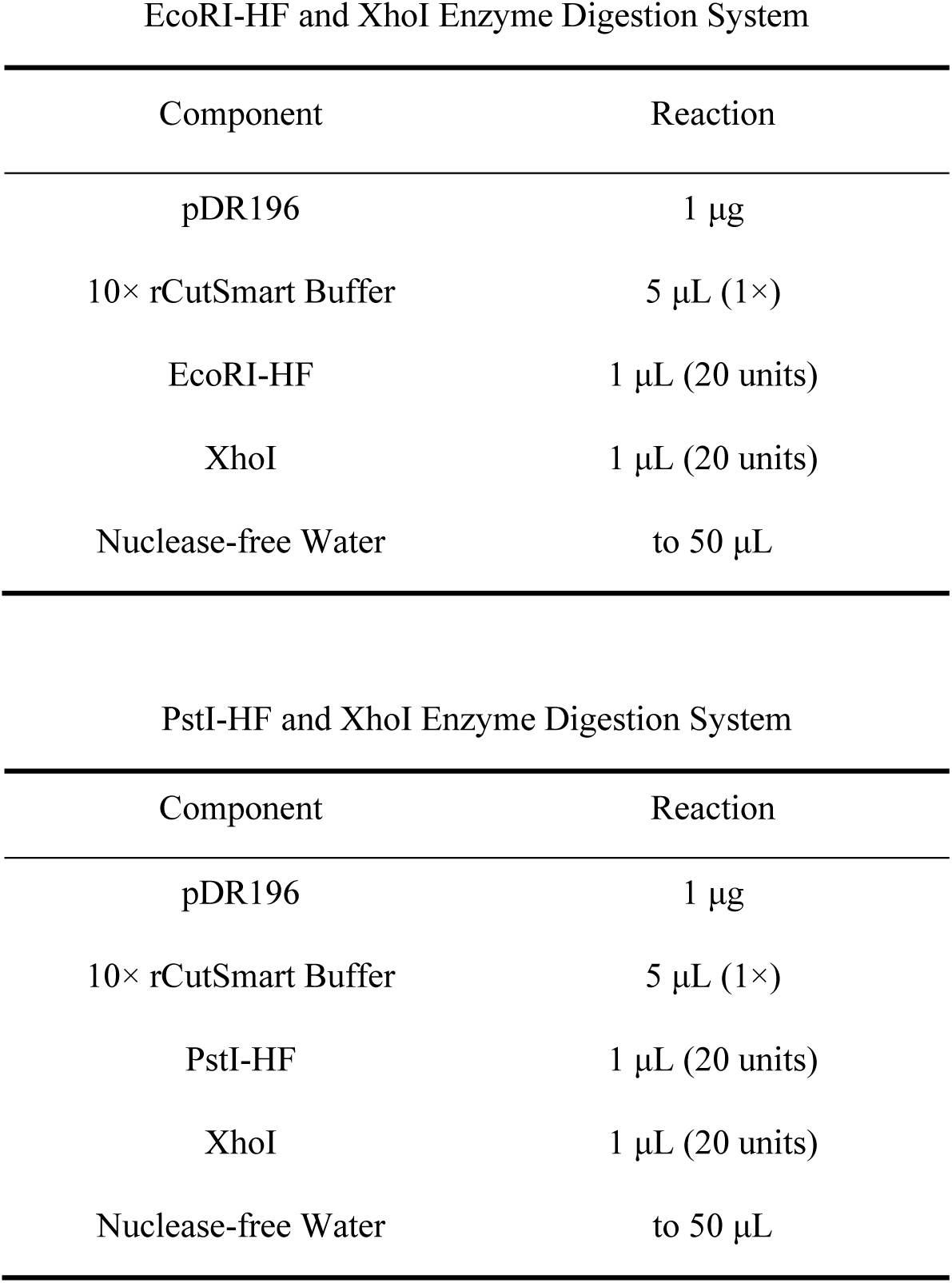
The enzyme digestion system utilized for the linearization of the pDR196 yeast expression vector

**Supplemental Table S5:**
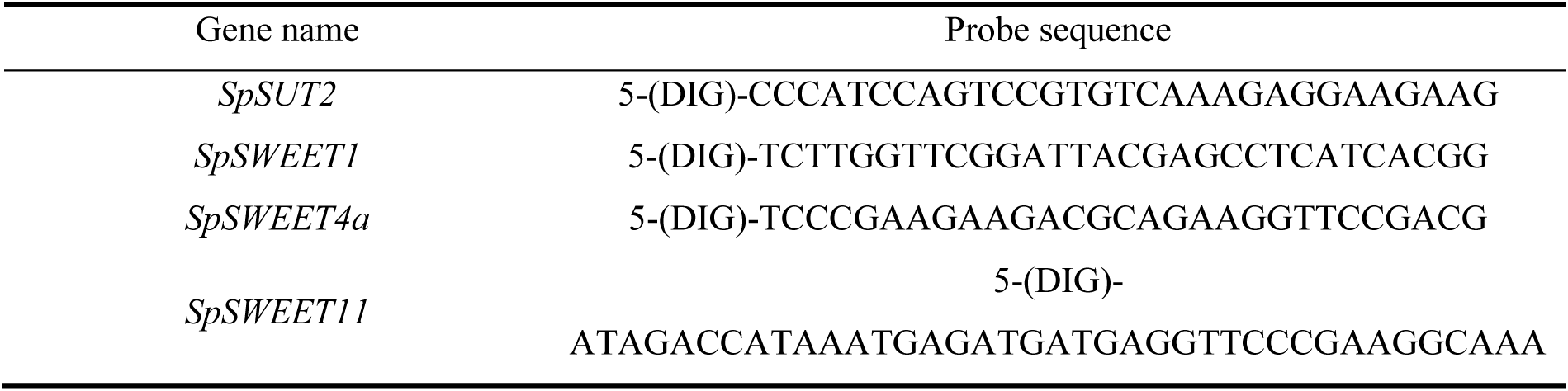
mRNA probe sequences for in situ hybridization

